# Controlling for confounding effects in single cell RNA sequencing studies using both control and target genes

**DOI:** 10.1101/045070

**Authors:** Mengjie Chen, Xiang Zhou

**Affiliations:** Departments of Biostatistics and Genetics, University of North Carolina, Chapel Hill, NC 27599.; Department of Biostatistics, Center for Statistical Genetics, University of Michigan, Ann Arbor, MI 48109..

**Keywords:** single cell RNA sequencing, partial least squares, confounding factors

## Abstract

Single cell RNA sequencing (scRNAseq) technique is becoming increasingly popular for unbiased and high-resolutional transcriptome analysis of heterogeneous cell populations. Despite its many advantages, scRNAseq, like any other genomic sequencing technique, is susceptible to the influence of confounding effects. Controlling for confounding effects in scRNAseq data is a crucial step for proper data normalization and accurate downstream analysis. Several recent methodological studies have demonstrated the use of control genes for controlling for confounding effects in scRNAseq studies; the control genes are used to infer the confounding effects, which are then used to normalize target genes of primary interest. However, these methods can be suboptimal as they ignore the rich information contained in the target genes. Here, we develop an alternative statistical method, which we refer to as scPLS, for more accurate inference of confounding effects. Our method is based on partial least squares and models control and target genes jointly to better infer and control for confounding effects. To accompany our method, we also develop a new,block-wise expectation maximization algorithm for scalable inference. Our algorithm is an order of magnitude faster than standard ones, making scPLS applicable to hundreds of cells and hundreds of thousands of genes. With extensive simulations and comparisons with other methods, we demonstrate the effectiveness of scPLS. Finally, we apply scPLS to analyze two scRNAseq data sets to illustrate its benefits in removing technical confounding effects as well as for removing cell cycle effects.

## 1. Introduction

Single-cell RNA sequencing (scRNAseq) has emerged as a powerful tool in genomics. While the traditional RNA sequencing, known as the bulk RNAseq, measures gene expression levels averaged across many different cells in a sample of potentially heterogeneous cell population, scRNAseq can measure gene expression levels directly at the single cell resolution. As a result, scRNAseq is less influenced by the variation of cell type and cell composition across different samples – a major confounding in the analyses of bulk RNAseq studies. Because of this benefit and its high resolution, scRNAseq provides unprecedented insights into many basic biological questions that are previously difficult to address. For example, scRNAseq has been applied to classify novel cell subtypes [49, 55] and cellular states [16, 31], reconstruct cell lineage and quantify progressive gene expression during development [47, 46, 9, 53], perform spatial mapping and re-localization [1, 39], identify differentially expressed genes and gene expression modulars [41, 21, 27], and investigate the genetic basis of gene expression variation by detecting heterogenic allelic specific expressions [4, 8].

Like any other genomic sequencing experiment, scRNAseq studies are influenced by many factors that can introduce unwanted variation in the sequencing data and confound the down-stream analysis [43]. Due to low capture efficiency and low amount of input material, such unwanted variation are exacerbated in scRNAseq experiments [50]. Indeed, adjusting for confounding factors in scRNAseq data has been shown to be crucial for accurate estimation of gene expression levels and successful down-stream analysis [13, 20, 22, 43, 50]. However, depending on the source, adjusting for confounding factors in scRNAseq can be non-trivial. Some confounding effects, such as read sampling noise and drop-out events, are direct consequences of low sequencing-depth, which are random in nature and can be readily addressed by probabilistic modeling using existing statistical methods [20, 13, 22, 10, 36]. Other confounding effects are inherent to a particular experimental protocol and can cause amplification bias, but can be easily mitigated by using new protocols [14]. Yet other confounding effects are due to observable batches and can be adjusted for by including batch labels and technician ids as covariates or dealt with other statistical methods [18, 51]. However, many confounding factors are hidden and are difficult or even impossible to measure. Common hidden confounding factors include various technical artifacts during library preparation and sequencing, and unwanted biological confounders such as cell cycle status. These hidden confounding factors can cause systematic bias, are notoriously difficult to control for, and are the focus of the present study.

To effectively infer and control for hidden confounding factors in scRNAseq studies, we develop a novel statistical method, which we refer to as scPLS. scPLS is specifically designed in the unsupervised setting where the predictor variables are not known *a priori* (e.g. cell clustering problems). scPLS takes advantage of the fact that genes in a scRNAseq study can often be naturally classified into two sets: a control set of genes that are free of effects of the predictor variables and a target set of genes that are of primary interest. By modeling the two sets of genes jointly using the partial least squares regression, scPLS is capable of making full use of the data to improve the inference of confounding factors. scPLS is closely related to and bridges between two existing subcategories of methods: a subcategory of methods (e.g. PCA [37, 28, 34, 44] and LMM [17, 19, 29]) that treat control and target genes in the same fashion, and another subcategory of methods (e.g. RUV [37, 15] and scLVM [6]) that use control genes alone for inferring confounding factors. By bridging between the two subcategories of methods, scPLS enjoys robust performance across a range of application scenarios. scPLS is also computationally efficient: with a new block-wise expectation maximization (EM) algorithm, it is scalable to thousands of cells and tens of thousands of genes. Using simulations and two real data applications, we show how scPLS can be used to remove confounding effects and enable accurate down-stream analysis in scRNAseq studies. Our method is implemented as a part of the Citrus project and is freely available at: http://chenmengjie.github.io/Citrus/.

The paper is organized as follows. In Section 2 we provide a brief review of existing statistical methods for removing confounding effects and describe how scPLS is related to and motivated from these methods. In Section 3, we provide a methodological description of the scPLS model, with inference details provided in Section 4. In Section 5 we present comparisons between scPLS and several existing methods using simulations. In Section 6, we apply scPLS to two real scRNAseq data sets to remove technical confounding effects or cell cycle effects. Finally, we conclude the paper with a summary and discussion in Section 7.

## 2. Review of Previous Methods

Many statistical methods have been developed in sequencing- and array-based genomic studies to infer hidden confounding factors and control for hidden confounding effects. Based on their targeted application, these statistical methods can be generally classified into two categories.

The first category of methods are supervised and application-specific: these methods are designed to infer the confounding factors in the presence of a *known* predictor variable, and to correct for the confounding effects without removing the effects of the predictor variable. For example, scientists are often interested in identifying genes that are differentially expressed between two pre-determined treatment conditions or that are associated with a measured predictor variable of interest. To remove the confounding effects in these applications, methods, include SVA [28], sparse regression models [45, 54], and, more recently, RUV [12, 11], are developed. Although these application-specific methods are widely applied in many genomics studies, their usage is naturally restricted to cases where the primary variable of interest is known. The application-specific methods become inconvenient in cases where there are multiple variables of interest (e.g. in eQTL mapping problems). They also become inapplicable when the primary variable of interest is not observed (e.g. in clustering problems).

Our scPLS belongs to the second category of unsurpervised methods, which are designed to infer the confounding factors without knowing or using the predictor variable of interest. Notable applications of unsurpervised methods in scRNAseq studies include cell type clustering and classification [49, 55, 16, 31, 47, 46, 9, 53]. Existing unsurpervised statistical methods can be further classified into two subcategories. The first subcategory of methods treat all genes in the same fashion and use all of them to infer the confounding factors. For example, the principal component analysis (PCA) or the factor model extracts the principal components or factors from all genes as surrogates for the confounding factors [37, 28, 34, 44]. The inferred factors are treated as covariates whose effects are further removed from gene expression levels before downstream analyses. Similarly, the linear mixed models (LMMs) construct a sample relatedness matrix based on all genes to capture the influence of the confounding factors [17, 19, 29]. The relatedness matrix are then included in the downstream analyses to control for the confounding effects. In contrast, the second subcategory of unsupervised methods are recently developed to take advantage of a set of control genes for inferring the confounding factors [6, 37]. These methods divide genes into two sets: a control set of genes that are known to be free of effects of interest *a priori* and a target set of genes that are of primary interest. Unlike the first subcategory, the second subcategory of methods treat the two gene sets differently in inferring the confounding factors: the confounding factors are only inferred from the control set, and are then used to remove the confounding effects in the target genes for subsequent downstream analysis. For example, scRNAseq studies often add ERCC spike-in controls prior to the PCR amplification and sequencing steps. The spike-in controls can be used to capture the hidden confounding technical factors associated with the experimental procedures, which are further used to remove technical confounding effects (e.g. reverse transcription or PCR amplification confounding effects) from the target genes [17]. Similarly, most scRNAseq studies include a set of control genes that are known to have varying expression levels across cell cycles. These cell cycle genes can be used to capture the unmeasured cell cycle status of each cell, which are further used to remove cell cycle effects in the target genes [6]. Prominent methods in the second subcategory include the unsupervised version of RUV [37, 15] and scLVM [6].

The two subcategories of unsupervised methods use different strategies to infer the confounding factors. Therefore, these two sets of methods are expected to perform well in different settings. Specifically, the first subcategory of methods have the advantage of using information contained in all genes to accurately infer the confounding effects. However, when the predictor variable of interest influences a large number of genes, then this subcategory of methods may incorrectly remove the primary effects of interest. On the other hand, the second subcategory of methods infer confounding factors only from the control genes and are thus not prone to mistakenly removing the primary effects of interest. However, these methods overlook one important fact – that the hidden confounding factors not only influence the control genes but also the target genes, i.e. the exact reason that we need to remove such confounding effects in the first place. Because the confounding factors influence both control and target genes, using control genes alone to infer the confounding factors can be suboptimal as it fails to use the information from target genes.

To more effectively infer and control for hidden confounding factors in scRNAseq studies, we develop a novel statistical method, which we refer to as scPLS. scPLS bridges between the two subcategories of unsupervised methods and effectively includes each as a special case. Like the first subcategory of methods, scPLS models both control and target genes jointly to infer the confounding factors. Like the second subcategory of methods, scPLS is capable of taking advantage of a control set to guild the inference of confounding factors. scPLS builds upon the partial least squares regression model and relies on a key modeling assumption that only target genes contain the primary effects of interest or other systematic biological variations. By incorporating such systematic variations in the target genes only, we can jointly model both control and target genes to infer the confounding effects while avoiding mis-removing the primary effects of interest. Therefore, scPLS has the potential to make full use of the data to improve the inference of confounding factors and the removal of confounding effects.

## 3. Details of scPLS

We provide modeling details for scPLS here. Our scPLS is generally applicable to both sequencing- and array-based genomic studies, but we focus on its application in scRNAseq. The scRNAseq data resembles that of the bulk RNAseq data and consists of a gene expression matrix on *n* cells and *p* + *q* genes. We consider dividing the genes into two sets: a control set that contains *q* control genes and a target set that contains *p* genes of primary interest. The control genes are selected based on the purpose of the analysis. For example, the control set would contain ERCC spike-ins if we want to remove technical confounding factors, and would contain cell cycle genes if we want to remove cell cycle effects. We use the following partial least squares regression to jointly model both control and target genes:

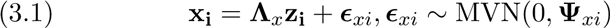

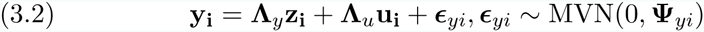

where for *i*’th individual cell, **x**_**i**_ is a *q*-vector of expression levels for *q* control genes; **y**_**i**_ is a *p*-vector of expression levels for *p* target genes; **z**_**i**_ is *k*_*z*_-vector of unknown confounding factors that affect both control and target genes; the coefficients of the confounding factors are represented by the *q* by *k*_*z*_ loading matrix **Λ**_*x*_ for the control genes and the *p* by *k*_*z*_ loading matrix **Λ**_*y*_ for the target genes; **u**_**i**_ is a *k*_*u*_-vector of unknown factors in the target genes and potentially represents the predictors of interest or other structured variations (see below); **Λ**u is a *p* by *k*_*u*_ loading matrix; Exi is a *q*-vector of idiosyncratic error with covariance 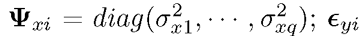 is a vector of idiosyncratic error with covariance 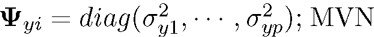 denotes the multivariate normal distribution. We assume **z**_**i**_~ MVN(0, **I**) and **u**_**i**_~ MVN(0, **I**). We model transformed data instead of the raw read counts. We also assume that the expression levels of each gene have been centered to have mean zero, which allows us to ignore the intercept.

scPLS includes two types of unknown latent factors. The first set of factors, **z**_**i**_, represents the unknown confounding factors that affect both control and target genes. The effects of **z**_**i**_ on the control and target genes are captured in the loading matrices **Λ**_*x*_ and **Λ**_*y*_, respectively. We call **z**_**i**_ the confounding factors throughout the text, and we aim to remove the confounding effects **Λ**_*y*_ **z**_**i**_ from the target genes. The second set of factors, **u**_**i**_, aims to capture a low dimensional structure of the expression level of *p* target genes. The factors **u**_**i**_ can represent the unknown predictor variables of interest, specific experimental perturbations, gene signatures or other intermediate factors that coordinately regulate a set of genes. Therefore, the factors **u**_**i**_ can be interpreted as cell subtypes, treatment status, transcription factors or regulators of biological pathways in different studies [7, 35, 30, 3, 33]. Although **u**_**i**_ could be of direct biological interest in many data sets, we do not explicitly examine the inferred **u**_**i**_ here. Rather, we view modeling **u**_**i**_ in the target genes as a way to better capture the complex variance structure there and to facilitate the precise estimation of confounding factors **z**_**i**_. For simplicity, we call **u**_**i**_ the biological factors throughout the text, though we note that **u**_**i**_ could well represent non-biological processes such as treatment or environmental effects. Thus, the expression levels of the control genes can be described by a linear combination of the confounding factors **z**_**i**_ and residual errors; the expression levels of the target genes can be described by a linear combination of the confounding factors **z**_**i**_, the biological factors **u**_**i**_ and residual errors. For both types of confounding factors, we are interested in inferring the factor effects **Λ**_*y*_ **z**_**i**_ and **Λ**u**u**_**i**_ rather than the individual factors **z**_**i**_ and **u**_**i**_. Therefore, unlike in standard factor models, we are not concerned with the identifiability of the factors. Figure 1 shows an illustration of scPLS.

**FIG 1.**
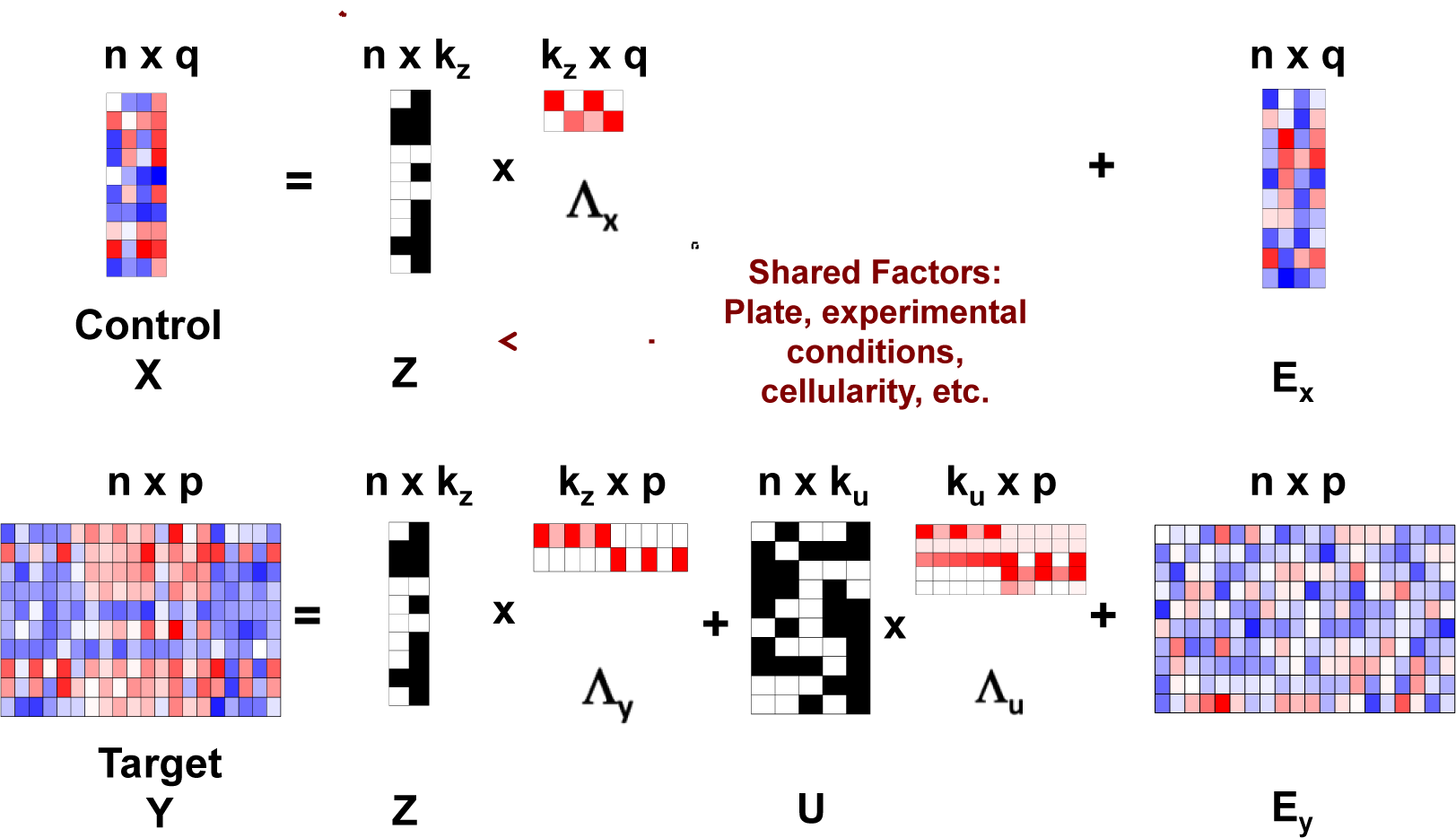
Illustration of scPLS. We model the expression level of genes in the control set (**X**) and genes in the target set (**Y**) jointly. Both control and target genes are affected by the common confounding factors (**Z**) with effects**Λ**x and **Λ**y in the two gene sets, respectively. The target genes are also influenced by biological factors (**U**) with effects **Λ**u. The biological factors represent intermediate factors that coordinately regulate a set of genes, and are introduced to better capture the complex variance structure in the target genes. **E**_**x**_ and **E**_**y**_ represent residual errors. scPLS aims to remove the confounding effects **ZΛ**_y_ in the target genes.

scPLS is closely related to the two subcategories of unsupervised methods described in Section 2. Specifically, without the biological effects term **Λ_u_u_i_**, scPLS effectively reduces to the first subcategory of methods that treat all genes in the same fashion for inferring the confounding factors. Without the Equation 3.2 term, scPLS effectively reduces to the second subcategory of methods that use only control genes for inference. (Note that, after inferring the confounding factors **z**_**i**_ from Equation 3.1, the second subcategory of methods still use a reduced version of Equation 3.2 without the biological effects term **Λ_u_u_i_** to remove the confounding effects.) By including both modeling terms, scPLS can robustly control for confounding effects across a range of scenarios. Therefore, scPLS provides a flexible modeling framework that effectively includes the two subcategories of unsupervised methods as special cases and has the potential to outperform these previous methods.

## 4. EM Algorithms for scPLS

We develop an expectation-maximization (EM) algorithm for inference in scPLS. Specifically, we first initialize the factor loading matrices (**Λ**_*x*_, **Λ**_*y*_, **Λ_u_**) based on sequential single value decompositions on the gene expression matrices (**X** = (**x**_**1**_, …, **x**_**q**_), **Y** = (**y**_**1**_, …, **y**_**p**_)) (Algorithm 1). Afterwards, we treat the latent factors (**w**_**i**_ = 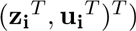 as missing data, use an iterative procedure to compute the expectation of the factors conditional on each individual cell data (**v**_**i**_ = 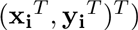 in turn in the E-step, and then update the factor loading matrices (**Λ** = (**Λ**_*x*_, **Λ**_*y*_, **Λ**_**u**_)) by merging information across all individuals in the M-step (Algorithm 2). We list the EM algorithm below, with detailed derivation provided in Appendix A.

### Algorithm 1 Initializer of EM algorithms for scPLS

**Input:** Data matrices **X**, **Y**, and the number of latent factors *k*_*z*_ and *k*_*u*_.

**Output: Λ**^(0)^, the initial value for **Λ**.

Apply SVD on **X**, obtain **U**, **D**, **V**;

Calculate 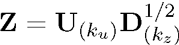 and standardize the elements in **Z** to have mean 0 and variance 1;

Use least squares to estimate 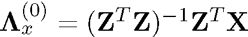 and 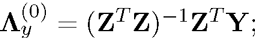

Obtain the residuals of **X** after removing the confounding effects, or 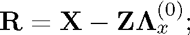

Similarly, apply SVD on **R**, obtain **U´**, **D´**, **V´**;

Calculate 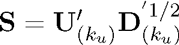 and standardize elements in **S** so that all elements have mean 0 and variance 1;

Use least squares to estimate 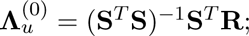

### Algorithm 2 Naive EM algorithm for scPLS

**Input:** Data **w**.

**Output:** 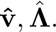

Initialize **Λ**^(0)^ using Algorithm 1;

Initialize ϋ^(0)^ = **I**;

**E step**: Compute *E*(**v**_**i**_*|***w**_**i**_)^(t)^ and *E*(**v**_**i**_**v**_**i**_T *|***w**_**i**_)^(t)^, given **Λ**(t), Λ(t);

**M step**:

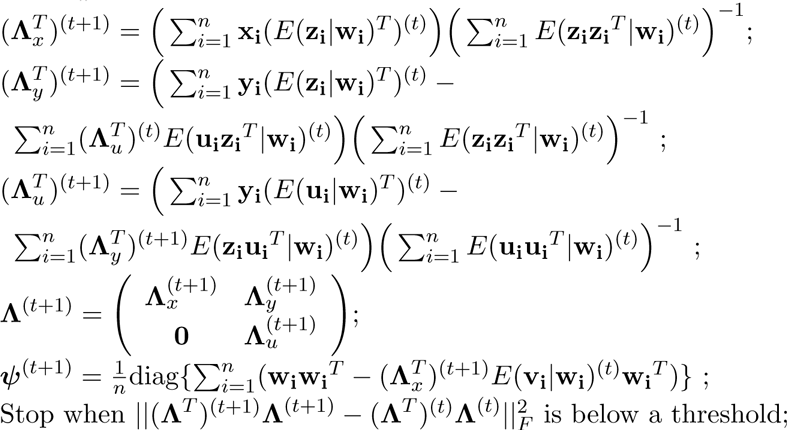

We refer to the above algorithm (Algorithm 2) as the naive EM algorithm. The naive EM algorithm is computationally expensive: it scales quadratically with the number of genes and linearly with the number of cells/samples. To improve the computational speed, we develop a new EM-in-chunks algorithm (Algorithm 3). Our algorithm is based on the observation that the expression levels of the target genes are determined by the same set of underlying factors and that these factors can be estimated accurately even with a small subset set of target genes. This allows us to randomly divide target genes into dozens of chunks, compute the expectation of the factors in each chunk separately in the E-step, and then average these expectations across chunks. With the averaged expectations, we then update the factor loading matrices in the M-step. Thus, our new algorithm modifies the E-step in the naive algorithm and becomes *K* times faster than the naive one, where *K* is the number of chunks. Simulations (detailed in Section 5) show that our EM-in-chunks algorithm yields almost comparable results to the naive EM algorithm with respect to estimation errors, but can be close to an order of magnitude faster (Table 1). Therefore, we apply the EM-in-chunks algorithm with chunk size 500 throughout the rest of the paper.

### Algorithm 3 EM-in-chunks algorithm for scPLS

**Input:** Data *W*.

**Output:** 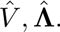

Initialize **Λ**^(0)^ using Algorithm 2; Initialize *Ψ^(0)^* = **I**;

Initialize *E*(**v**_**i**_*|***w**_**i**_)^(0)^ and *E*(**v**_**i**_**v**_**i**_T *|***w**_**i**_)^(0)^ using E step in Algorithm 1;

**M step**:

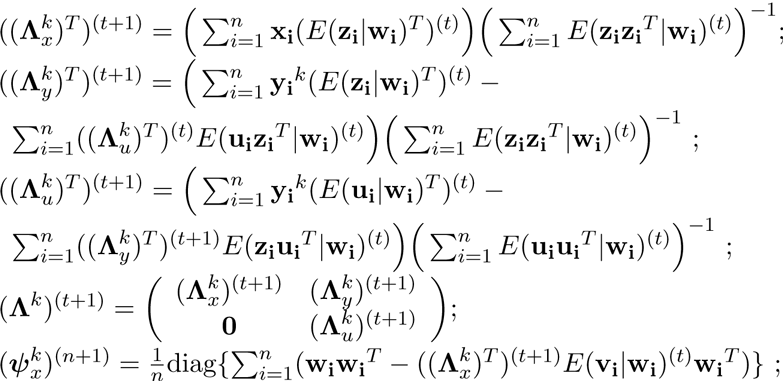

**E step**:

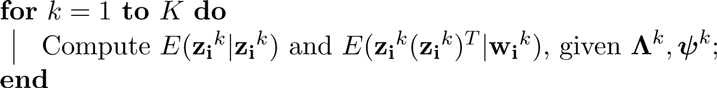

Average among *K* chunks and obtain 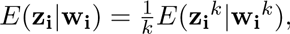

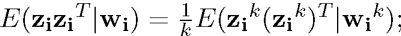

Iterate between M and E step until last cycle;

Given *E*(**z**_**i**_*|***w**_**i**_) and *E*(**z**_**i**_**z**_**i**_^*T*^ *|***w**_**i**_) from the last cycle, the final estimate of iΛ and Ψ are calculated using one M step in Algorithm 1;

Finally, we use the Bayesian information criterion (BIC) to determine the number of confounding factors *k*_*z*_ and the number of biological factors *k*_*u*_. Specifically, we evaluate the likelihood on a grid of *k*_*z*_ (1 to 3) and *k*_*u*_ values (1 to 10) and choose the optimal combination that minimizes the BIC. After estimating the model parameters on the optimal set of *k*_*z*_ and *k*_*u*_, we use the residuals 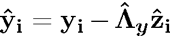 as the de-noised values for subsequent analysis.

Note that the residuals are only free of the confounding effects **Λ**_*y*_ **z**_**i**_ but still contain the biological effects **Λ**_**u**_**u**_**i**_.

## 5. Simulations

We performed a simulation study to compare scPLS with other methods. Specifically, we simulated gene expression levels for 50 control genes and 1,000 target genes for 200 cells. These 200 cells come from two equal-sized groups, representing two treatment conditions or two sub-cell types. Among the 1,000 target genes, only 100 of them are differentially expressed (DE) between the two groups and thus represent the signature of the two groups. The effect sizes of the DE genes were simulated from a normal distribution, and we scaled the effects further so that the group label explains a fixed percentage of phenotypic variation (PVE) in expression levels in the DE genes (ranging from 1% to 20%, with 1% increment). In addition to the group effects, we set *k*_*z*_ = 2*, k*u = 5 and simulated each element of **z**_**i**_ and **u**_**i**_ from a standard normal distribution. We simulated each element of Λ_*x*_ from *N*(-0.25,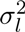) and each element of **Λ**_*y*_ from *N* (0.25, 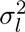Note that **Λ**_*x*_ and **Λ**_*y*_ were simulated differently to capture the fact that the effect sizes of the confounding factors could be different for control and target genes. We simulated each element of **Λ**_**u**_ from *N* (0, 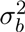). We simulated each element of *ɛ*_*xi*_ and *ɛ*_*yi*_ from a standard normal distribution. We set 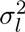 = 0.4 and 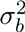 = 0.6 to ensure that, in non-DE genes, the confounding factors **z**_**i**_ explain 20% PVE in either the control or the target genes; the biological factors **u**_**i**_ explain 30% PVE of the target genes; and the residual errors to explain the rest of PVE. After we simulated gene expression levels, we further converted these continuous values into count data by using a Poisson distribution: the final observation for *i*th cell and *j*th gene *c*ij is from *c*_*ij*_ *~* Poi(*N* exp(*μ* + *w*_*ij*_)), with *w*_*ij*_ being the continuous gene expression levels simulated above and *N* = 500000, *=* = log(10/500000). Note that, because of the residual errors, the resulting count data are over-dispersed with respect to a Poisson distribution.

**Table 1.**
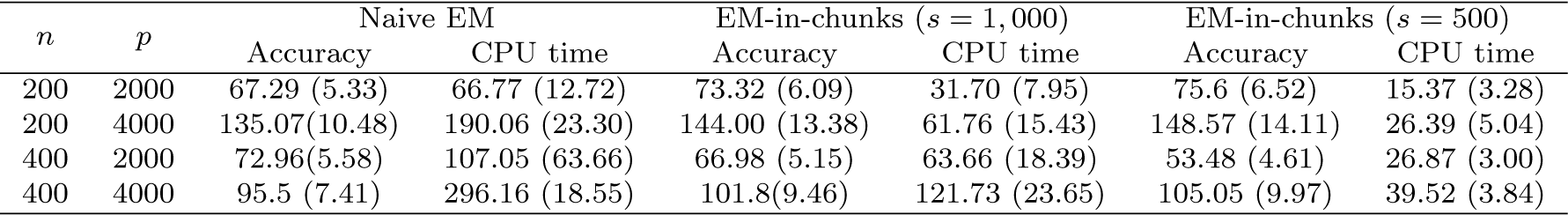
Comparison of the naive EM algorithm and the EM-in-chunks algorithm in terms of accuracy and speed. The EM-in-chunks algorithm uses either a chunk size of 500 genes or a chunk size of 1,000 genes. Accuracy is measured by the estimation error of the loading matrix in terms of the normalized Frobenius norm 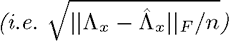. Speed is measured by CPU time in minutes. Standard deviations across 10 replicates are listed inside parenthesis. s: number of genes per chunk. n: the number of cells. p: the number of genes in the target set. The number of genes in the control set is q = 50 in all simulations.

| $n$ | $p$ | Naive EM | | EM-in-chunks ($s = 1,000$) | | EM-in-chunks ($s = 500$) | |
| --- | --- | --- | --- | --- | --- | --- | --- |
|  |  | Accuracy | CPU time | Accuracy | CPU time | Accuracy | CPU time |
| 200 | 2000 | 67.29 (5.33) | 66.77 (12.72) | 73.32 (6.09) | 31.70 (7.95) | 75.6 (6.52) | 15.37 (3.28) |
| 200 | 4000 | 135.07(10.48) | 190.06 (23.30) | 144.00 (13.38) | 61.76 (15.43) | 148.57 (14.11) | 26.39 (5.04) |
| 400 | 2000 | 72.96(5.58) | 107.05 (63.66) | 66.98 (5.15) | 63.66 (18.39) | 53.48 (4.61) | 26.87 (3.00) |
| 400 | 4000 | 95.5 (7.41) | 296.16 (18.55) | 101.8(9.46) | 121.73 (23.65) | 105.05 (9.97) | 39.52 (3.84) |

We considered three different simulation scenarios. In scenario I, the con-founding factors **z**_**i**_ are independent of group labels. In scenario II, the con-founding factors are correlated with group labels. To simulate correlated data, we simulated each element of **z**_**i**_ from *N* (0,1) if the corresponding sample belongs to the first group, but from *N* (-0.25,1) if the corresponding sample belongs to the second group. Finally, we also considered a scenario III where there is no biological factor (i.e. data were simulated effectively under the PCA modeling assumption and all genes could be used to infer the confounding factors). We performed 10 simulation replicates for each scenario.

We compared our method to four different methods: (1) PCA and (2) LMM (implemented in GEMMA [56, 57]) all genes used to infer the confounding effects; while (3) RUVseq (version 1.2.0); which we simply refer to as RUV in the following text) and (4) scLVM (version 0.99.1) only control genes used to infer the confounding effects. We used default settings in all the above methods. We used the count data directly for RUV and used log transformed data (i.e. log(*c*_*ij*_ + 1)) for all other methods. For PCA and RUV, we set the number of latent factors to be the true number (i.e. 2). Such number is determined automatically by the software itself for scLVM, and is not needed for LMM. Our goal on the simulated data is twofold: we want to identify these differentially expressed genes and to classify the 200 cells into two groups. Therefore, we compared the performance of various methods based on two criteria: the power to identify the DE genes and the power to classify cells into two groups. We permuted group labels to construct an empirical null and compared methods based on either power given 5% false discovery rate (FDR) for identifying DE genes.

It is useful to point out that the three simulation scenarios are designed to highlight the hybrid nature of scPLS. In particular, in the presence of biological factors (i.e. scenarios I and II), the methods that use all genes to remove confounding effects, such as PCA and LMM, may incorrectly remove the primary effects of interest. Therefore, we would expect RUV and scLVM to outperform PCA and LMM in scenarios I and II. In the absence of biological factors (i.e. scenarios III), the methods that only use control genes to remove confounding effects, such as RUV and scLVM, may fail to utilize all information contained in the data. Therefore, we would expect PCA and LMM to outperform RUV and scLVM in scenario III. Because of the hybrid nature of scPLS, we would expect it to perform well across all scenarios.

Simulation results confirm our expectations. Specifically, in scenario I (Figure 2a), scPLS outperforms the other four methods in identifying DE genes across a range of PVEs. Among the rest of the four methods, RUV and scLVM outperform PCA and LMM. Similarly, in scenario II (Figure 2b),

**Fig 2.**
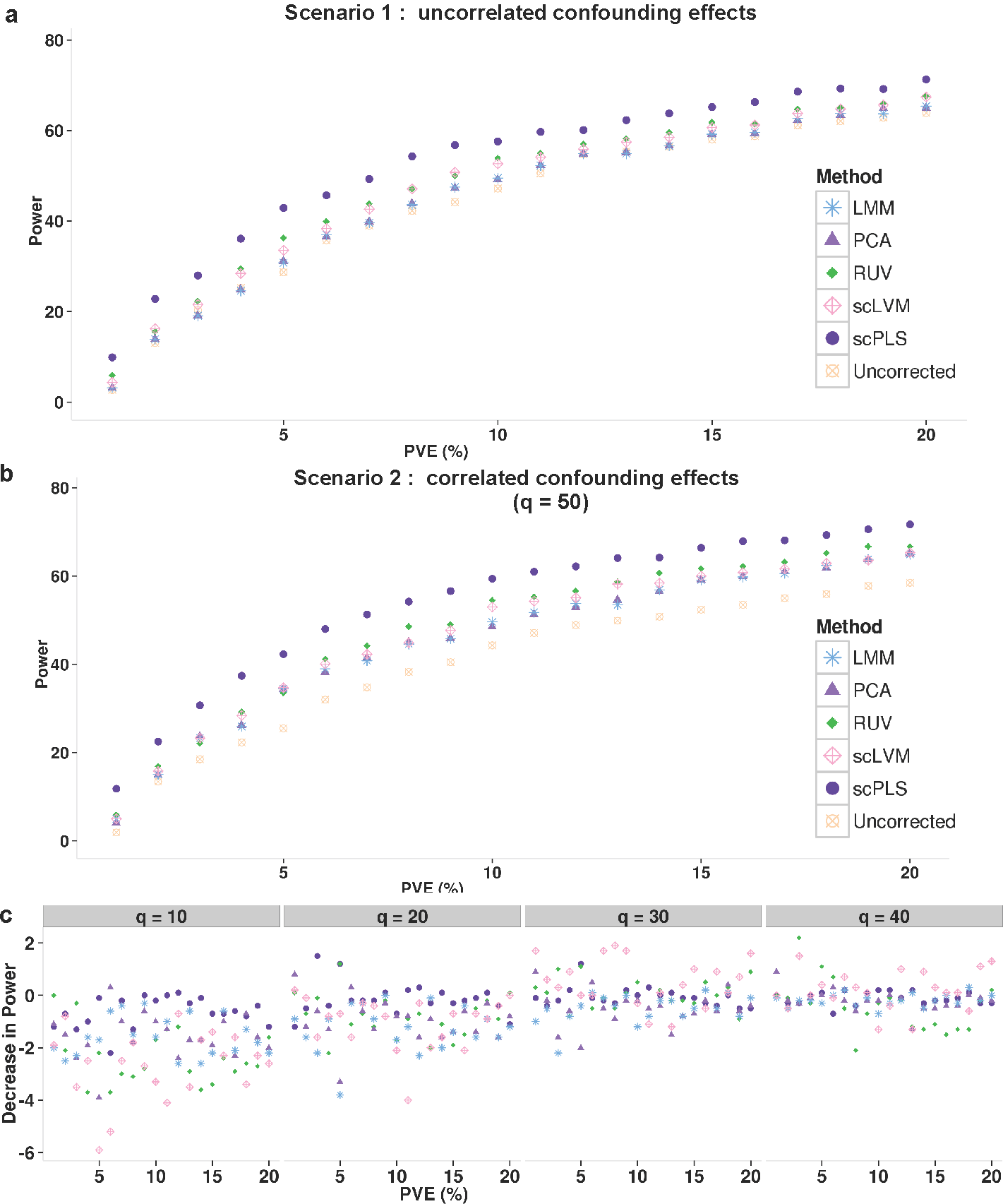
Method comparison in simulations. Identifying differentially expressed genes using scPLS-corrected data achieves higher power than using LMM-, PCA-, RUV- and scLVM- corrected data or uncorrected data in both scenario I (a) and scenario II (b) across a range of effect sizes. Power is evaluated at an empirical false discovery (FDR) rate of 0.05 and is averaged across ten simulation replicates. x-axis shows the effect sizes, which are measured as the percentage of phenotypic variation (PVE) in expression levels explained by the group label (ranges from 1% to 20%). (c) Sensitivity analysis shows that, compared with other methods, scPLS has the least percentage reduction in power (y-axis) when a smaller subset of control genes are used (q = 10, 20, 30 or 40 instead of 50).

scPLS performs the best, followed by RUV and scLVM. PCA and LMM perform the worst. Compared with RUV and scLVM, scPLS is also more robust with respect to the number of control genes used in the analysis (Figure 2c). In particular, because scPLS does not completely rely on the information contained in the control genes, it achieves good performance even if we only use a much smaller subset of control genes. In contrast, the performance of RUV and scLVM compromises more quickly when a reduced number of control genes is used (especially when using *q* = 10). The higher power of scPLS to detect DE genes in scenario I and II also translates to a better performance of classifying single cells (Figure 3a). To quantify the classification performance, we applied the support vector machine (SVM) to classify the cells. We performed a five-fold cross-validation, training SVM with 80% of the samples and evaluating the prediction accuracy with the rest of the samples. All methods achieve similarly high power in the easiest case when PVE is greater than 10%. However scPLS outperforms the other four methods when PVE is low and the classification task is difficult. For example, in scenario I, when PVE = 1%, scPLS achieves an average accuracy of 77% across 10 replicates, while LMM, PCA, RUV and scLVM achieve 71.2%, 71.2%, 73.5%, 70.5%, respectively.

**Fig 3.**
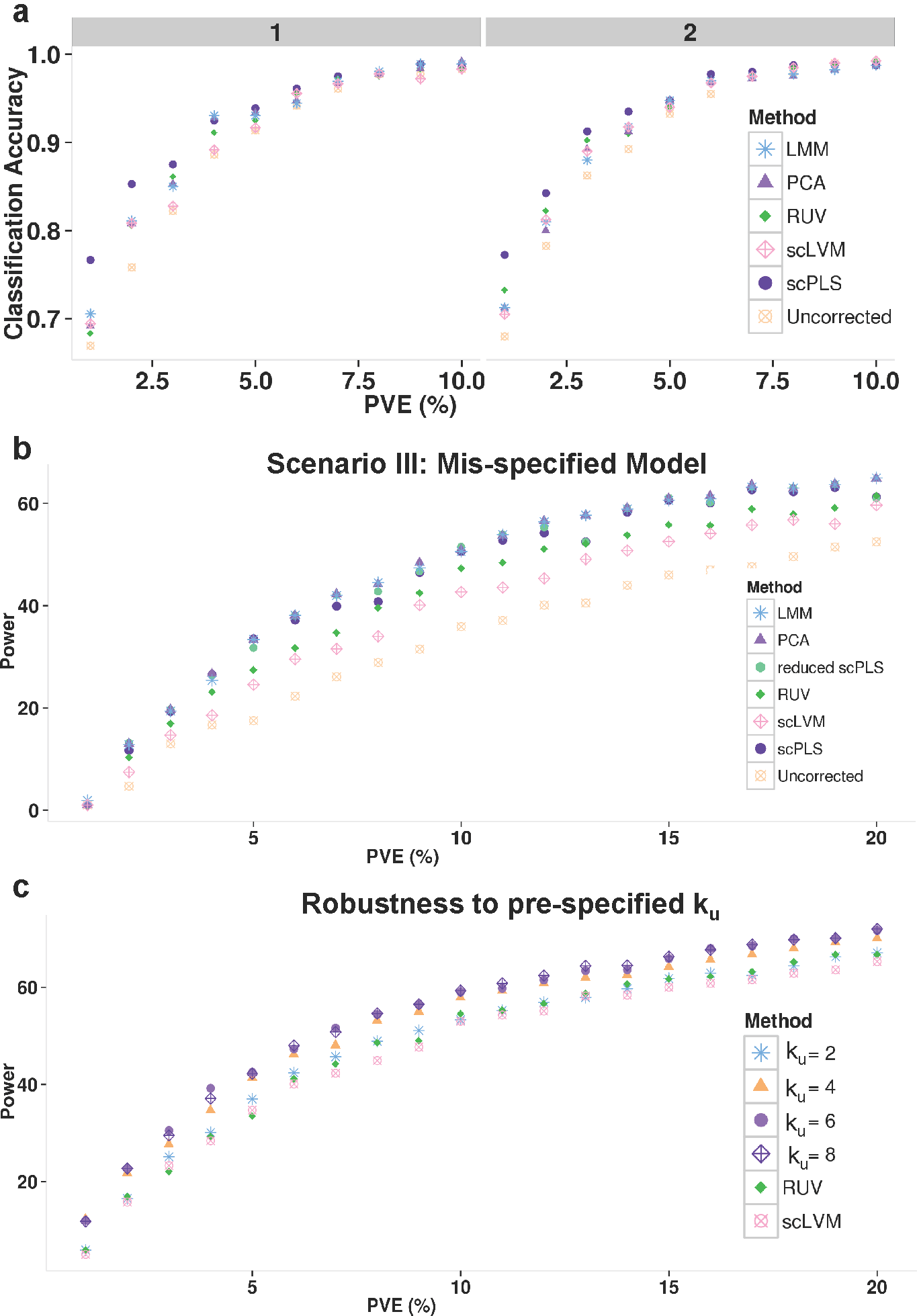
Method comparison in simulations (continued). (a) scPLS corrected expression data can be used to better classify cells into the two known clusters than LMM-, PCA-, RUV- and scLVM-corrected data or uncorrected data in both scenario I and scenario II across a range of effect sizes. Classification is based on support vector machine (SVM) with five-fold cross-validation. Accuracy is computed as the mean percentage of true positives in the test set across replicates. (b) Identifying differentially expressed genes using scPLS-corrected data achieves similar power as using LMM-, PCA-corrected data, all of which are more powerful than RUV- and scLVM-corrected or uncorrected data in scenario III across a range of effect sizes. (c) scPLS is robust with respect to ku, as the power to identify differentially expressed genes remains similar when a different number of biological factors is used (ku=2, 4, 6, 8) in scenario III where in turth ku = 0. x-axis shows the effect sizes, which are measured as the percentage of phenotypic variation (PVE) in expression levels explained by the group label (ranges from 1% to 20%).

On the other hand, in scenario III, scPLS performs as well as PCA and LMM, and all these three methods outperform RUV and scLVM (Figure 3b). Importantly, scPLS is not sensitive with respect to the number of biological factors used in fitting the model, and achieves similar power for a range of reasonable *k*_*u*_ values when the truth is 0 in scenario III (Figure 3c). As it is often unknown whether a low-rank structural variation exists in a real data set, our simulation suggests that we can always include the biological factors ***u***_**i**_ in the model even in the absence of such factors.

Therefore, the simulation results highlight the hybrid nature of scPLS.

scPLS works robustly well across a range of scenarios while the other two subcategories of methods work preferentially well only in scenarios that most favor their modeling assumptions.

## 6. Real Data Applications

Next, we applied scPLS to two real data sets. The first dataset is used to demonstrate the effectiveness of scPLS in removing the technical confounding effects by using ERCC spike-ins. Removing technical confounding effects is a common and important task in transcriptome analysis. The second dataset is used to demonstrate the effectiveness of scPLS in removing cell cycle effects by using a known set of cell cycle genes. Removing cell cycle effects can reveal gene expression heterogeneity that is otherwise obscured.

### 6.1. Removing Technical Confounding Factors

The first dataset consists of 119 mouse embryonic stem cells (mESCs), including 74 mESCs cultured in a two-inhibitor (2i) medium and 45 mESCs cultured in a serum medium [13]. We obtained the raw UMI counts data directly form the authors and the data contains measurements for 92 ERCC spike-ins and 23,459 genes. Due to the low coverage of this dataset (median coverage equals one), we filtered out lowly expressed genes and selected only genes that have at least five counts in more than a third of the cells. This filtering step resulted in a total of 17 ERCC spike-ins that were used as the controls and 2,772 genes that were used as the targets. As in the simulations, we log transformed the count data and centered the transformed values for scPLS, PCA, LMM and scLVM. We used the count data for RUV. In this data, scPLS infers *k*_*z*_ = 1 confounding factors and *k*_*u*_ = 1 biological factors. In the target genes, the confounding factors and structured biological factors explain a median of 20% and 19% of gene expression variance, respectively. The PVE by the confounding and biological factors can be as high as 86.2% and 76.9%, respectively, in the target genes.

We applied scPLS and the other four methods to remove confounding effects in the data. To compare the performance of different methods in the real data, we performed a clustering analysis. We reason that if method is effective in removing confounding effects, then the corrected data from the method could be used to separate the mESCs into the two known clusters (i.e. 2i medium vs serum medium). For the clustering analysis, we applied the k-means method, an unsupervised method, with the number of clusters set to two, on uncorrected data and data corrected by different methods. Consistent with our simulations, scPLS outperforms all other methods based on a variety of clustering performance measurements (Table 2).

### 6.2. Removing Cell Cycle Effects

Our method can also be used to remove cell cycle effects. To demonstrate its effectiveness there, we applied scPLS and several other methods to a second dataset [6]. This dataset contains gene expression measurements on 9,570 genes from 182 embryonic stem cells (ESCs) with pre-determined cell-cycle phases (G1, S and G2M). The uncorrected data we obtained are already pre-processed by the original study to remove the technical effects and are thus continuous. Therefore, we did not apply RUV here. To remove cell cycle effects, we used 629 annotated cell-cycle genes as controls and the other genes as targets. scPLS infers *k*_*z*_ = 1 cell cycle confounding factors, and *k*_*u*_ = 1 biological factors. These factors explain a median of 0.4% and 0.1% of gene expression variance, respectively. The PVE by cell cycle factors and biological factors can be as high as 7% and 2%, respectively. We visualized the uncorrected data and scPLS corrected data on a PCA plot (Figure 4). In the uncorrected data, there is a clear separation of cells according to cell-cycle stage. Such separation of cells is not observed in the corrected data, indicating that the cell cycle related expression signature is effectively removed.

**Table 2.**
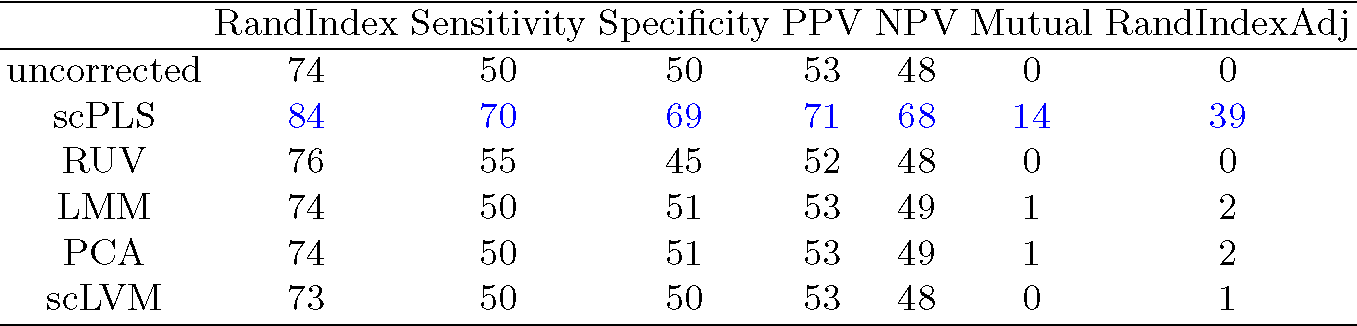
A number of clustering measurements show that the corrected data by scPLS can be used to better reveal the two known clusters than the other four methods in the first data set. A k-means algorithm (with two clusters) is applied to the uncorrected data and data corrected by different methods. Clustering performance is measured by Rand Index, sensitivity, specificity, positive predictive value (PPV), negative predictive value (NPV), mutual information (Mutual) or adjusted Rand Index (RandIndexAdj). Blue color labels the best performer by each criterion. All performance measurements are averaged across 10 runs and are multiplied by a factor of 100.

We compared scPLS and the other three methods in their effectiveness in removing cell cycle effects. Following the original study [6], we evaluated method performance based on the following criteria. Specifically, we computed for each gene the proportion of expression variance explained by the cell cycle factor. We denote this quantity as PVEi, which stands for inferred PVE. Because the cell-cycle stage of each cell had been experimentally determined in this data set, we further computed the variance explained by the true cell cycle labels. We denote this quantity as PVEt, which stands for true PVE. For scPLS, PVEi and PVEt are highly correlated (*r*^*2*^ = 0.94), demonstrating the efficacy of scPLS. The correlation remains the same whether we use the full control set or with a subset of 300 controls. The correlation between PVEi and PVEt in scPLS is slightly higher, with statistical significance, than scLVM (*r*^*2*^ = 0.92; p-value < 10^−16^ comparing scPLS vs scLVM), LMM (*r*^*2*^ = 0.92; p-value < 10^−16^ comparing scPLS vs LMM), and PCA (*r*^*2*^ = 0.92; p-value < 10^−16^ comparing scPLS vs PCA). In addition, as an alternative measurement, the median of the absolute difference between PVEi and PVEt across genes from scPLS, scLVM, LMM and PCA are 0.018, 0.023, 0.019 and 0.019, respectively, again supporting a small advantage of scPLS. Therefore, the results suggest that scPLS works slightly better than the other three methods, though all methods work reasonably well in removing cell cycle effects in this data set (which is consistent with the low variance explained by the confounding factors).

**Fig 4.**
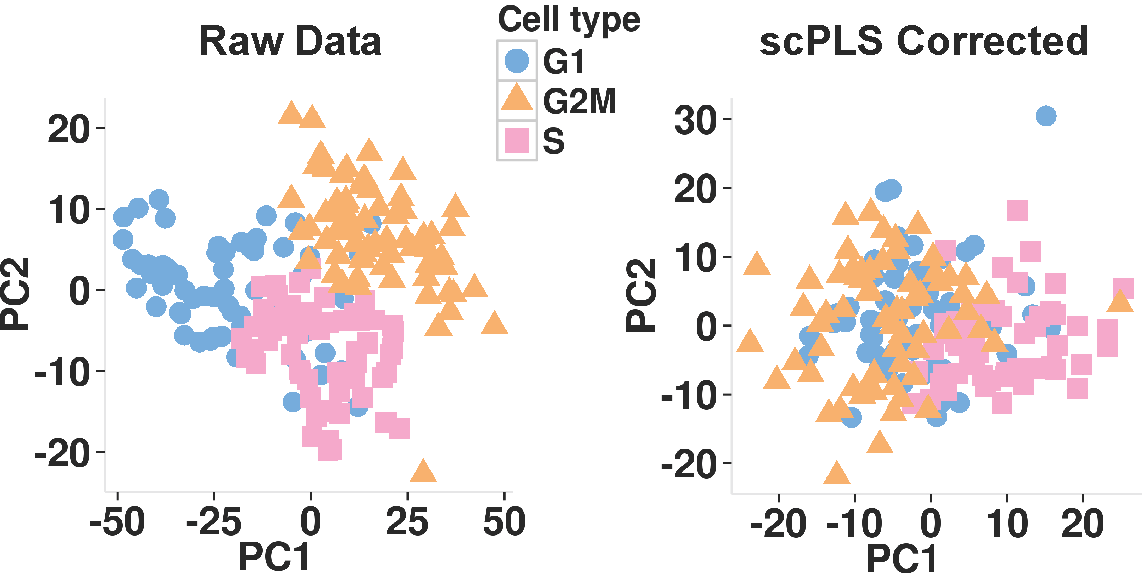
*PCA plots for the uncorrected data and scPLS corrected data in the second dataset. In the uncorrected data, there is a clear separation of cells by cell-cycle stage. Such separation of cells is no longer observed in the scPLS corrected data.*

## 7. Discussion

We have presented scPLS for removing hidden confounding effects in scRNAseq studies. scPLS models both control and target genes jointly to infer the confounding factors and shows robust performance across a range of application scenarios. With simulations and applications to two real data sets, we have demonstrated its effectiveness for removing technical confounding effects or cell cycle effects in scRNAseq studies.

Although we have focused on its applications to scRNAseq studies, scPLS can be readily applied to other genomic sequencing studies. For instance, our method can be used to remove confounding effects from gene expression levels in bulk RNAseq studies [48] or from methylation levels in bisulfite sequencing studies [25]. The main requirement of our method is a set of pre-specified control genes that are measured together with the target genes in the sequencing studies. It is often straightforward to obtain such control genes. For example, many scRNAseq studies include a set of ERCC spike-in controls that could be used to model and remove technical confounding effects [17]. Even when such ERCC spike-in controls are not present or when they are unreliable [37], we can select a known set of house-keeping genes as controls to remove technical confounding [37]. Similarly, we can use a set of known cell cycle genes to remove cell cycle effects. Importantly, the performance of scPLS is robust to the number of genes included in the control set and yields comparable results even when a much smaller number of control genes is used. This is because scPLS not only uses information from control genes but also relies on information from target genes. Insensitivity to the control set makes scPLS especially suited to removing confounding factors in studies where a control set is not clearly defined. Because of its effectiveness and robustness, we expect scPLS to be useful in removing confounding effects in a wide variety of sequencing studies.

One important feature of scPLS is that it includes a low-rank component to model the structured biological variation often observed in real data. By decomposing the (residual) gene expression variation into a low-rank structured component that is likely to be contributed by a sparse set of biological factors, and an unstructured component that reflects the remaining variation, scPLS can better model the residual error structure for accurate inference of confounding effects. Although here we have focused on using the biological factors to better infer the confounding effects, we note that the low-rank biology factors themselves could be of direct interest. In fact, low-rank factors inferred from many data sets using standard factor models have been linked to important biological pathways or transcription factors [7, 35, 30, 3, 33]. Inferring the biological factors using scPLS is not feasible at the moment, however: because of model identifiability, scPLS can only be used to infer the biological effects (i.e. **Λ**_**u**_**u**_**i**_) but not the biological factors (i.e. **u**_**i**_). That said, additional assumptions can be made on the structure of the factors or the factor loading matrices to make factor inference possible [52]. For example, we could impose sparsity assumptions on the low-rank factors to facilitate the inference of a parsimonious set of biological factors. Exploring the use of biological factors in scPLS is an interesting avenue for future research.

Like many other methods for scRNAseq [5] or bulk [24, 38] RNAseq studies, scPLS requires a data transformation step that converts the count data into quantitative expression data. Different transformation methods can affect the interpretation of the data and are advantageous in different situations [43]. Because scPLS does not rely on a particular transformation procedure, scPLS can be paired with any transformation methods to take advantage of their benefits. One potential disadvantage of scPLS is that it does not model raw count data directly. However, despite the count nature of sequencing data, it has been show that there is often a limited advantage of modeling the raw read counts directly, at least for RNAseq studies [42, 40]. Statistical methods that convert and model the quantitative expression data have been shown to be robust [24, 38] and most large scale bulk RNAseq studies in recent years have used transformed data instead of count data [23, 34, 2, 32]. However, we note that, unlike bulk RNAseq studies, single cell RNAseq data often come with low read depth. In low read depth cases, modeling count data while accounting for over-dispersion or dropout events in single cell RNAseq studies may have added benefits [20, 50]. Therefore, extending our framework to modeling count data [26, 58] is another promising avenue for future research.

## Appendix A Em Algorithms for Scpls

To derive the EM algorithm, we first integrate out the latent variables **z**_**i**_ and **u**_**i**_

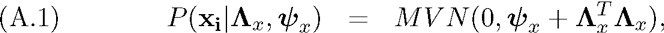

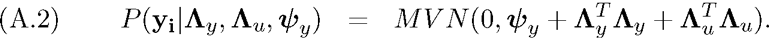

The latent variable **y**_**i**_ and **z**_**i**_ follow a joint normal distribution

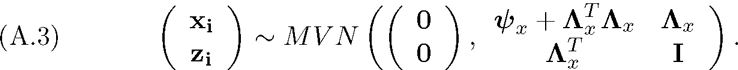

Denoting 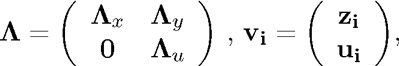 and 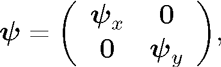 we can re-write 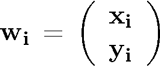 as **w**_**i**_ = Λ ^*T*^v_i_ + ψ. The variables **w**_**i**_ and **v**_**i**_ then follow a joint normal distribution

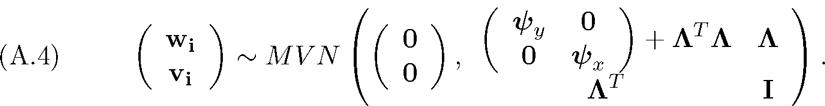

We view the latent factors **v**_**i**_ as the missing data. In the E step, we calculate the expectation of the log likelihood function for complete data. The expectation is taken with respect to the conditional distribution of **v**_**i**_ given **w**_**i**_

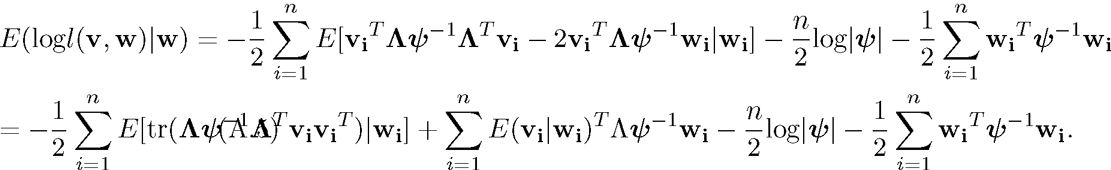

In the M step, we maximize the above expectation. To do so, we take derivatives of the log-likelihood function with respect to **Λ**_*x*_, **Λ**_*y*_ and **Λ**_**u**_, and obtain

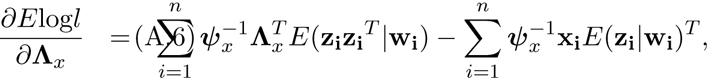

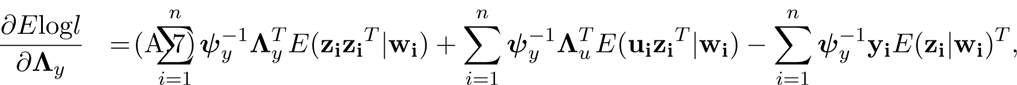

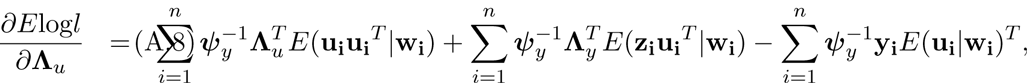

where the conditional expectations are

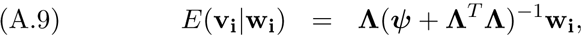

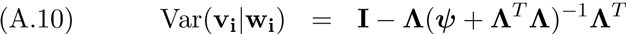

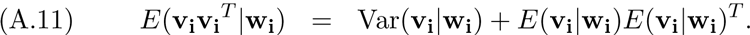

The above equations form the basis of our EM algorithms.

## ACKNOWLEDGEMENTS

MC was supported by the National Institutes of Health (NIH) grants R01 GM105785, R01 CA082659 and P01 CA142538. XZ was supported by NIH grants R01HL117626 (PI Abecasis) and R21ES024834 (PI Pierce), and a grant from the Foundation for the National Institutes of Health through the Accelerating Medicines Partnership (BOEH15AMP, co-PIs Boehnke and Abecasis). We thank Dr. Dominic Grun for providing the raw read counts of the first dataset.

